# Identification of biomarkers associated with ovarian cancer based on transcriptome sequencing

**DOI:** 10.1101/728618

**Authors:** jiazhou chen, xiandong peng, min yu

**Affiliations:** Obstetrics and Gynecology Hospital of Fudan University

**Keywords:** ovarian cancer, transcriptome sequencing, gene, pathway

## Abstract

**Objective:** This study aimed to explore more biomarkers associated with ovarian cancer.

**Methods:** Cell lines SKOV-3 (ovarian serous carcinoma cells) and MCV152 (benign ovarian epithelial tumor cell) were used in this study and performed transcriptome sequencing. The differentially expressed genes (DEGs) between ovarian cancer cells (SKOV-3) and controls (MCV152) were identified, followed by function enrichment analysis. The expression levels of genes involved in the key pathway were validated through PCR and western blot analyses.

**Results:** Total 2,020 upregulated and 1,673 downregulated DEGs were obtained between SKOV3 and MCV152 cells. The upregulated and downregulated DEGs were significantly associated with cell adhesion. In addition, the upregulated DEGs were significantly involved in pathways of ECM-receptor interaction, and the downregulated DEGs were involved in PI3K-Akt signaling pathway. PCR and western blot analyses showed that genes (proteins) expression related to PI3K-Akt signaling pathway were in consistent with bioinformatics analysis.

**Conclusion:** Cell adhesion and extracellular matrix (ECM)-receptor interaction may play an important role in the invasion of ovarian cancer. PI3K-Akt signaling pathway may be involved in the progression of ovarian cancer by up-regulating *ANGPT2*, *FGF18*, *ITGB4* and *ITGB8*, and downregulating *AKT3* and *PIK3AP1*.

**Highlights:**

1. Cell adhesion and ECM-receptor interaction may play important roles in ovarian cancer invasion.
2. PI3K-Akt signaling pathway may involve in ovarian cancer progression.
3. *ANGPT2*, *FGF18*, *ITGB4*, *ITGB8*, *AKT3* and *PIK3AP1* may serve as biomarkers in ovarian cancer.

## Introduction

Ovarian cancer is the seventh most common cancer among women worldwide as well as the tenth most common cancer in China [1]. It is estimated that there are 239,000 new cases of ovarian cancer and 152,000 deaths worldwide each year [2]. Despite improved treatment strategies, the overall survival rate of ovarian cancer is still as low as about 46%. According to the study, about 5% of female cancer deaths are due to low survival rates [3]. This mainly results from the intricate pathogenesis of ovarian cancer, making it difficult to find effective treatment and improve prognosis[4–6]. Therefore, a better understanding of the biological mechanisms of ovarian cancer is substantial for its earlier diagnosis and effective treatment.

Genetic variation has been determined to have functional effects on human cancers [7]. Molecular genetic analyses have revealed genetic alterations of several genes in ovarian cancers, such as *p53*, *c-erb-B2*, *c-myc* and *EGFR* [8, 9]. Previous microarray analysis has been applied to profile several genes, such as *SERPIN2*, *CD24* and *SEMACAP3*, that may serve as molecular markers for diagnosis and therapy of ovarian cancer [10]. In addition to genes, many pathways have also been demonstrated to involve in the progression of ovarian cancer. For instance, Wnt/β-catenin pathway could mediate the initiation of epithelial ovarian cancer cells by targeting genes that regulate cell proliferation and apoptosis [11]. The phosphatidylinositol 3 kinase (PI3K) pathway is found frequently altered in ovarian cancer and may serve as a therapeutic target [12]. Despite these findings, improving clinical outcomes of ovarian cancer are still far from enough. Therefore, there is an urgent need to explore more biomarkers.

In this study, we used two cell lines, SKOV-3 and MCV152, for transcriptome sequencing and identified the DEGs between ovarian cancer cells and normal controls. Subsequently we performed function analysis to investigate the potential pathways. The expression levels of genes involved in key pathway were validated at both mRNA and protein levels.

## Materials and methods

### Cell line and cell culture

Two cell lines were used in this study. MCV152 cell is benign ovarian epithelial tumor cell SV40-transformed serous papillary cystadenoma and transfected with telomerase hTERT. SKOV-3 cell is an ovarian serous carcinoma cell line purchased from the American Type Tissue Collection (Manassas, VA, USA). Both cell lines were cultured in Eagle’s minimal essential medium and maintained at 37°C in a humidified incubator containing 5% CO_2_.

### RNA extraction, library preparation and sequencing

RNA was extracted from cells using TRIzol® Reagent (Life Technologies, USA) according to the manufacturer’s protocol, and RIN scores were determined using an Agilent 2100 Bioanalyzer (Agilent Technologies, USA). One microgram RNA was used for cDNA library construction. The poly(A) mRNA was purified from total RNA using Dynal™ oligo(dT)-attached magnetic beads (Invitrogen, USA). The mRNA was cleaved into small fragments (∼ 200nt) for the synthesis of first strand cDNAs using random hexamer-primers and reverse transcriptase (SuperScript® II Reverse Transcriptase, Invitrogen). Second-strand cDNA was synthesized using DNA Polymerase I and RNase H (Invitrogen). The 3’ ends of DNA fragments were adenylated using Klenow DNA polymerase and amplified by PCR. cDNA libraries were sequenced using an Illumina HiSeq2000 sequencer.

### Quality control and genome mapping

The raw reads were cleaned by removing adaptor sequences, reads with > 2% undefined nucleotides (N), and low-quality reads containing > 20% of bases with qualities of < 20 using FastQC. Clean reads were aligned to hg38_UCSC_2013 genome using the star program [7].

### Data processing and differential expression analysis

Gene expression values were normalized using RPKM (reads per kilo base million reads) and FPKM (fragments per kilo base million reads), which were then performed principal component analysis (PCA). DESeq2 algorithm was applied to filter the DEGs. The significant criteria were: |log2 fold change (FC)| > 2 and false discovery rate (FDR) < 0.05. Heatmap plots were drawn by the R based on the differential expression analysis and the color was determined by the filtering criteria. Volcano plots were drawn by the R based on the DEGs analysis and the color was determined by the filtering criteria.

### Gene ontology (GO) and pathway analyses

GO analysis was performed to elucidate the biological implications of unique genes in the significant or representative profiles of the DEGs [8]. GO analysis included biological process (BP), molecular function (MF) and cellular component (CC) analyses. Pathway analysis was used to find out the significant pathway of the DEGs according to KEGG database. Fisher’s exact test was applied to identify the significant GO categories and pathways, and FDR was used to correct the p-values.

Additionally, the significant GO terms (*p<0.05*) were selected based on the up- and downregulated DEGs to construct the GO-Tree to summarize the function affected in this study [10]. The significant pathways (*p<0.05*) enriched by the up- and downregulated DEGs were selected to construct the Path-Act-Network using Cytoscape.

### Western blot

Total protein was extracted from cells using lysis buffer, followed by protein concentration determination. Protein sample (50 μL) was separated on 10% sodium dodecyl sulfate polyacrylamide gel electrophoresis (SDS-PAGE), and then transferred to polyvinylidenedifluoride (PVDF) membranes. The membrane was first incubated with primary antibodies (1:1000; ANGPT2, CD19, COL4A3, FGF18, ITGB4, ITGB8, LAMA3, LAMC2,PPP2R2C, SGK2, SYK, AKT3, COL6A1, CSF3, FGF1, ITGA2, ITGA11, MYB, PCK2, PGF, PIK3AP1, SGK1, TLR4 and TP53) at 4°C overnight and then incubated with the corresponding horseradish peroxidase (HRP)-conjugated secondary antibody (1:5000) for 1 h at 37°C. The blots were visualized using Image Lab software version 5.1 (Bio-Rad Laboratories, USA).

### Quantitative real-time PCR

Total RNA was extracted from cells using TRIzol Reagent (Sigma, USA). Complementary DNA (cDNA) was synthesized using a Reverse Transcription Kit (RR047A; Takara, Dalian, China). Real-time PCR was performed on a Light-Cycler Roche 480 (Roche Molecular Systems) with the Light-Cycler Roche 480 master kit. The results were normalized to the expression of GAPDH and calculated with the 2^−ΔΔCt^ method.

### Statistical analysis

All data were analyzed using GraphPad prism 5.0 software (GraphPad Prism, San Diego, CA). Results were expressed as mean ± standard deviation (SD). Differences for three or more groups were analyzed using one-way ANOVA tests. P value < 0.05 was considered to be statistically significant.

## Results

### Data processing and DEGs selection

A total of 46,499,161 clean reads were obtained after quality control and the mapping rate for sequencing data was 87.7%. PCA showed that the great sample correlation between MCV152 and SKOV3 cell lines (Figure 1). With thresholds of |log2 FC| > 2 and FDR < 0.05, a total of 3,693 DEGs were obtained, including 2,020 upregulated (such as *TACSTD2*, *KRT5*, *LINC00624*, *CDH16* and *PPP1R1B*) and 1,673 downregulated (such as *SNORA9*, *LINC00174*, *ZNF215*, *PPFIBP2* and *NTRK1*) ones, which are shown in heatmap (Figure 2A) and volcano plots (Figure 2B) based on the DEGs. The results above suggested that these DEGs could distinguish obviously in two samples.

**Figure 1.**
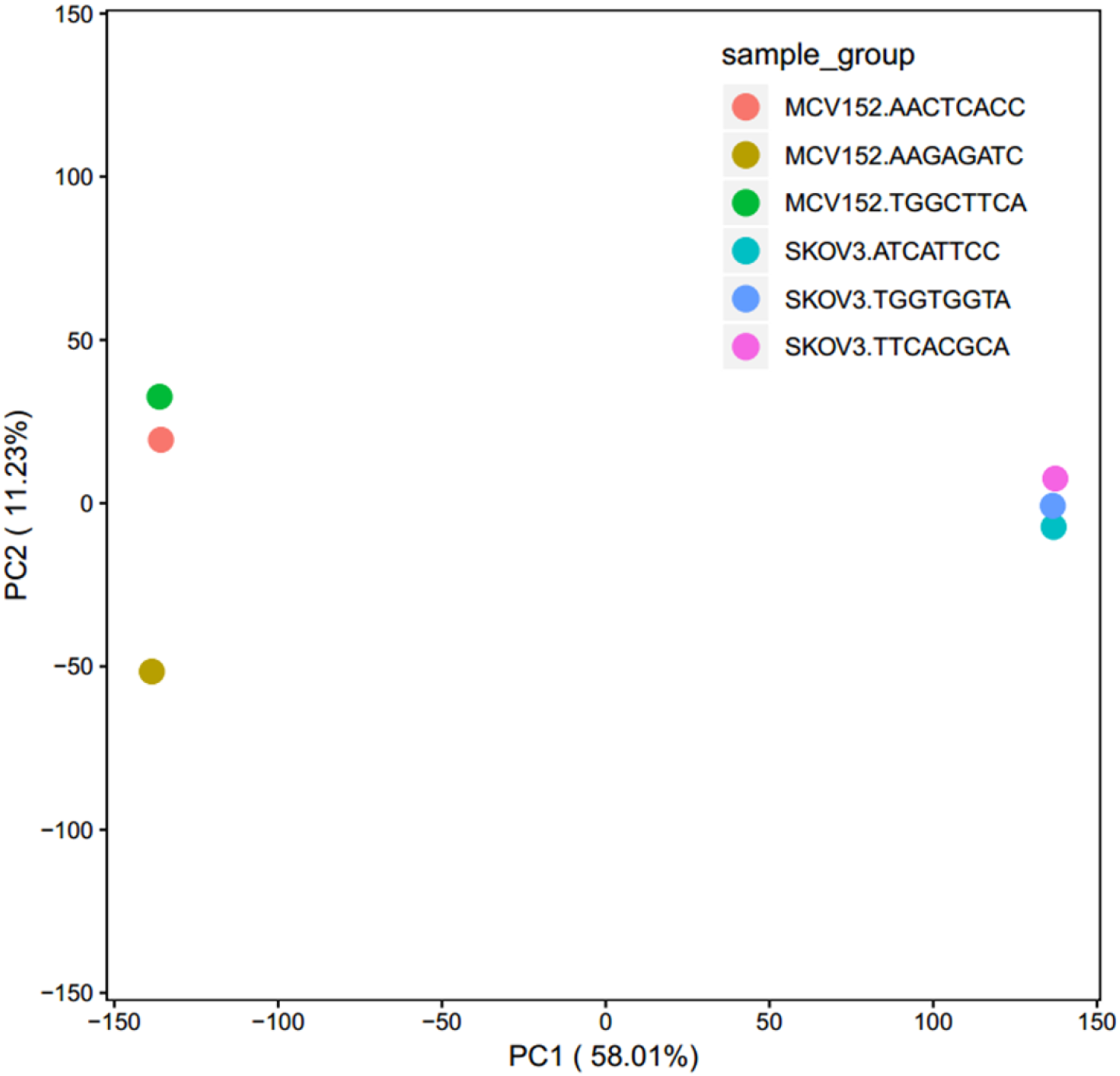
The mapping of all samples in a two-dimensional diagram composed of the largest component and the second largest component. The gene expression differences were small and correlation was good for RNA-seq samples with short distance, while the gene expression differences were large and the sample correlation was poor for RNA-seq samples with long distance. The gene expression difference of the shorter RNA sequence samples was smaller and the correlation was better, while the gene expression of the longer RNA sequence samples was different, and the sample correlation was poor.

**Figure 2.**
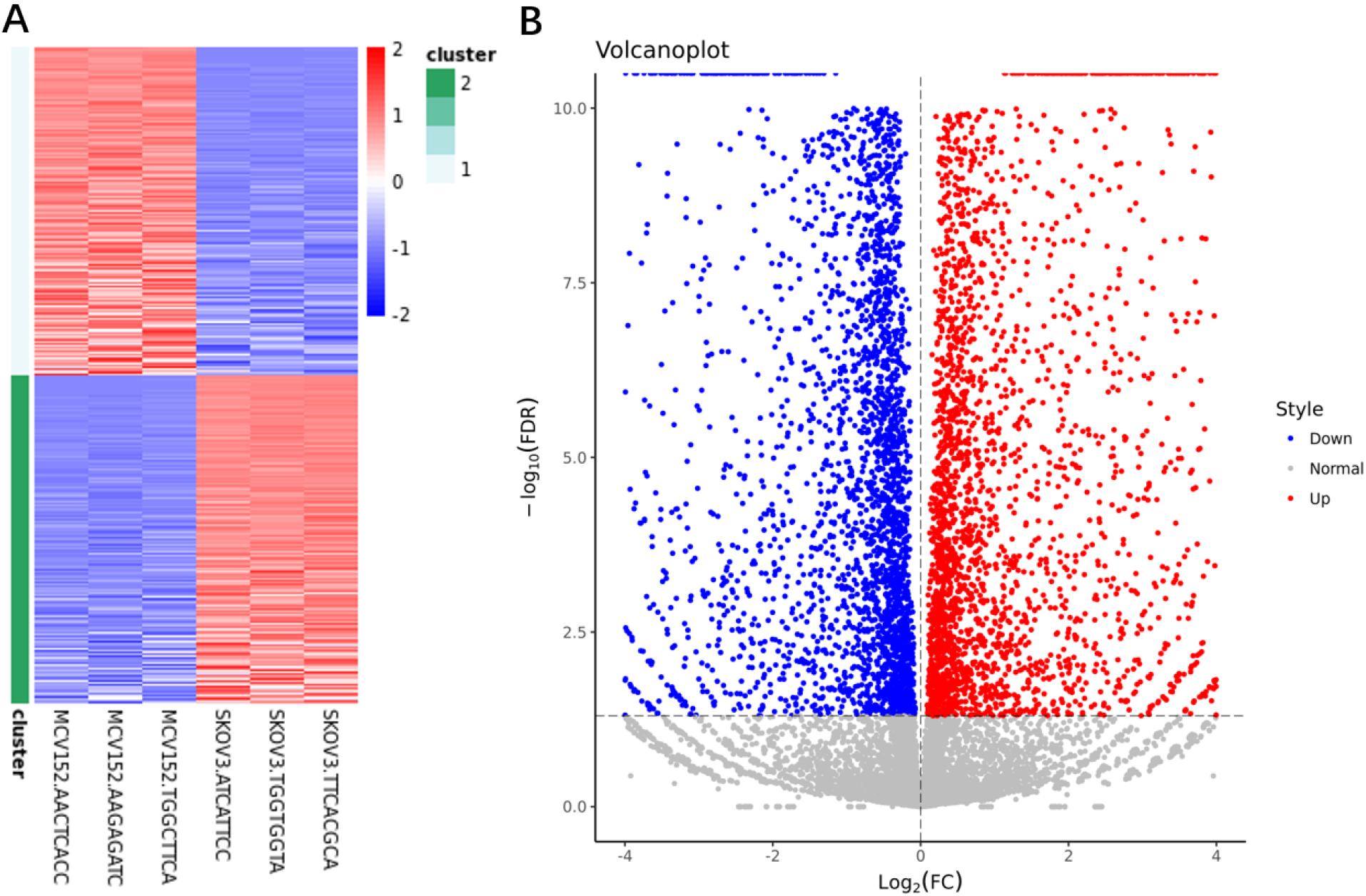
(A) Heatmap of differentially expressed genes. X axis: sample name; Y axis: gene name. (B) Volcano plot of differentially expressed genes. X axis: log2FC; Y axis: −log10 (FDR). red represents up-regulated gene, and blue represents down-regulated gene.

### GO and pathway analyses

The upregulated DEGs were significantly enriched in 401 BP terms, 134 MF terms and 66 CC terms. among which the top 15 terms for each category were present (Figure 3A). The BP terms were associated with cell adhesion, synapse assembly, and extracellular matrix organization; the MF terms were associated with calcium ion binding, and structural constituent of cytoskeleton; the CC terms were related to plasma membrane, and integral component of membrane. Additionally, the top 15 terms for each category are selected among the downregulated DEGs, which were significantly enriched in 665 BP terms, 139 MF terms and 82 CC terms (Figure 3B). For BP terms, the terms were associated with inflammatory response, immune response, and cell adhesion. The MF terms were related to receptor binding, and cytokine activity. The CC terms were associated with extracellular region, and plasma membrane. Moreover, pathway analysis showed that the upregulated genes were significantly involved in 33 pathways, such as extracellular matrix (ECM)-receptor interaction, axon guidance, and arrhythmogenic right ventricular cardiomyopathy (ARVC), and cell adhesion molecules (CAMs) (Figure 4A). The downregulated DEGs were significantly involved in 75 pathways, such as cytokine-cytokine receptor interaction, rheumatoid arthritis, pertussis, and PI3K-Akt signaling pathways (Figure 4B).

**Figure 3.**
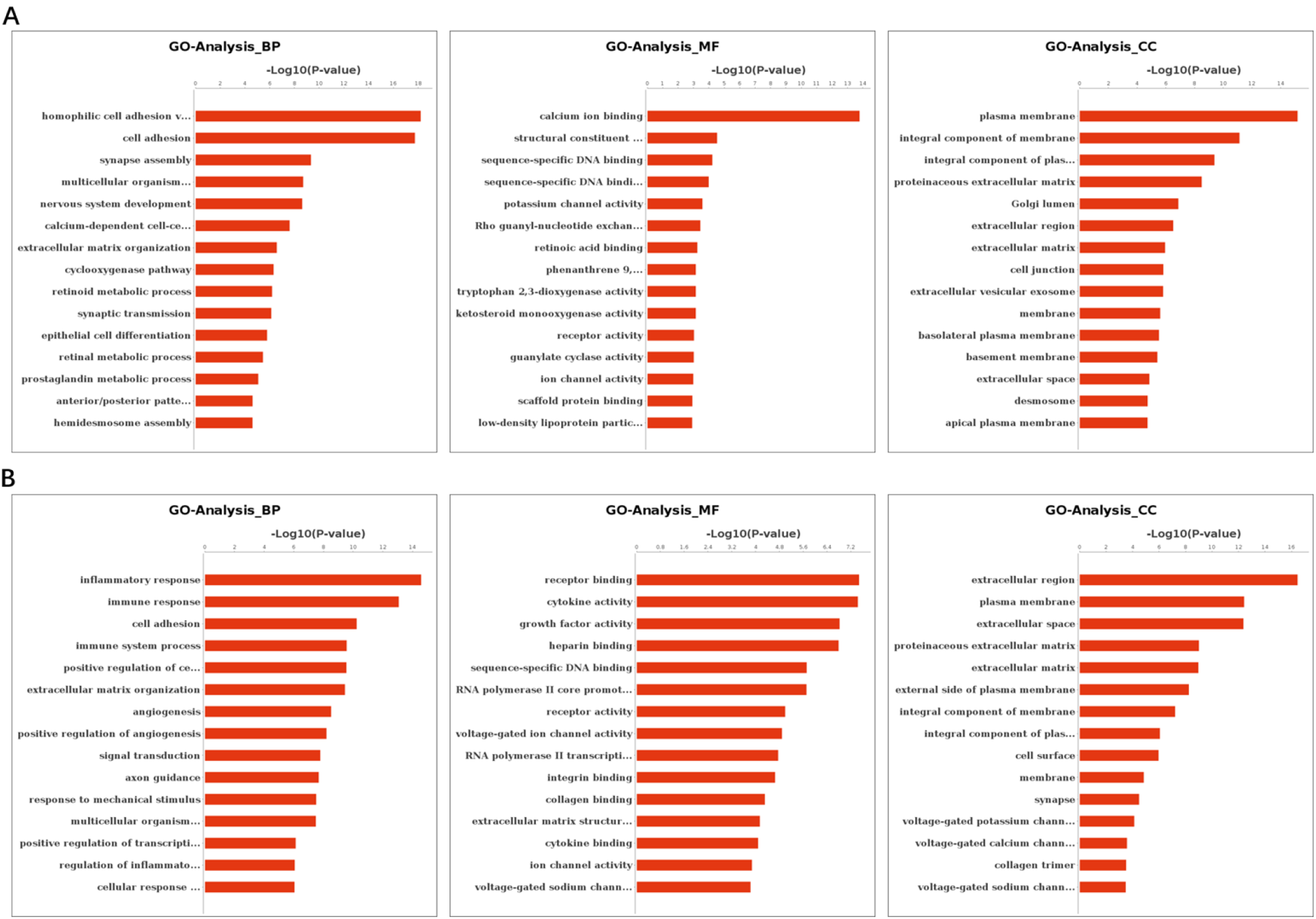
The GO functions (biological process (BP), molecular function (MF) and cellular component (CC)) enriched by upregulated genes (A) and downregulated genes (B).

**Figure 4.**
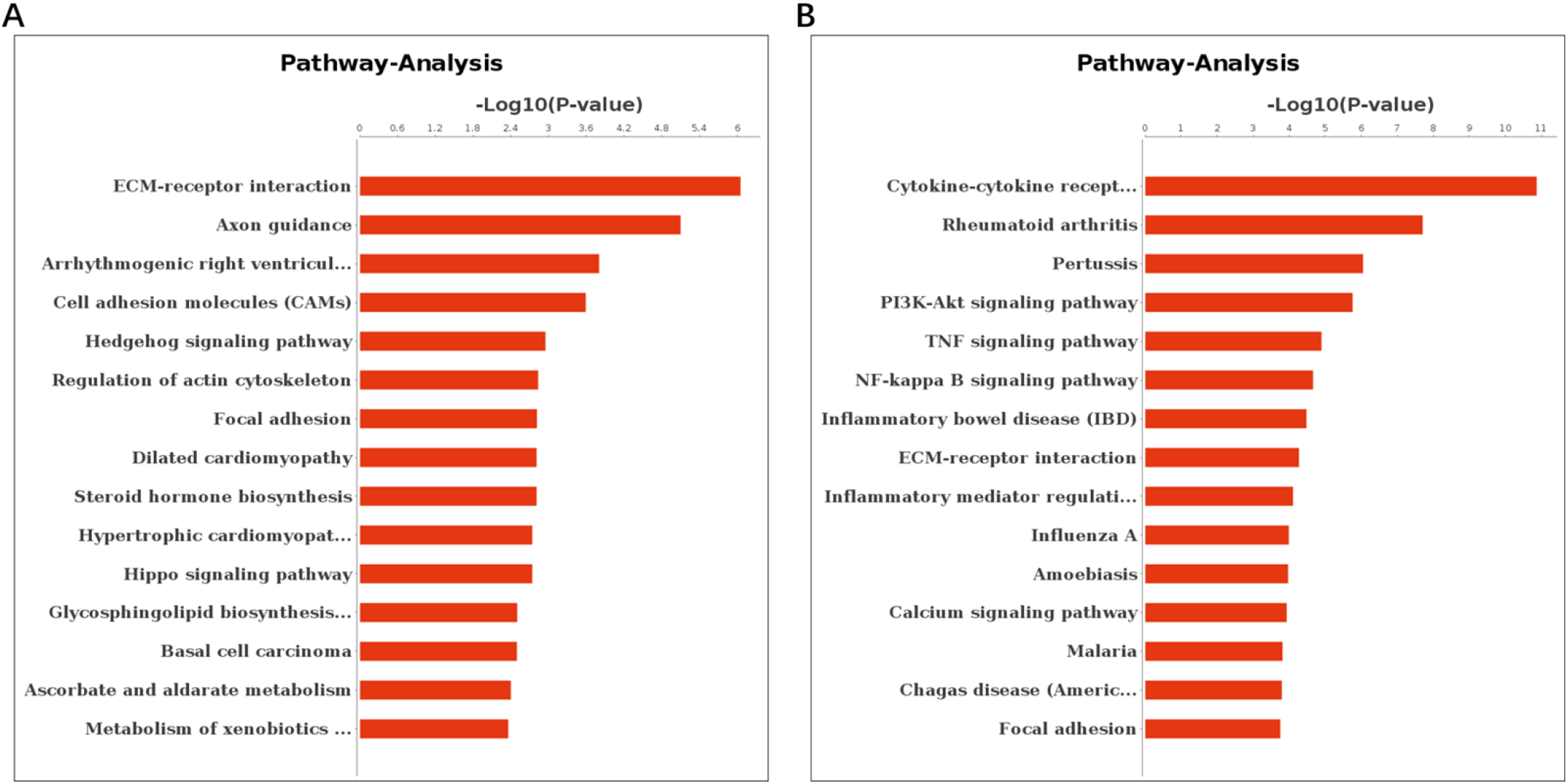
The pathways enriched by upregulated genes (A) and downregulated genes (B).

### GO and pathway network analyses

The GO-Tree and Path-Act-Network are shown in Figure 5. Cell migration had interactions with 6 GO terms, such as regulation of cell migration, positive regulation of cell migration, and axon guidance. Axon guidance also had interactions with 6 GO terms, such as motor neuron axon guidance, axon choice point recognition, and retinal ganglion cell axon guidance. Angiogenesis had interactions with 5 terms, like patterning of blood vessels, sprouting angiogenesis, and negative regulation of angiogenesis (Figure 5). PI3K-Akt signaling pathway interacted with 25 pathways, such as Ras signaling pathway, Rap1 signaling pathway, and p53 signaling pathway (Figure 6).

**Figure 5.**
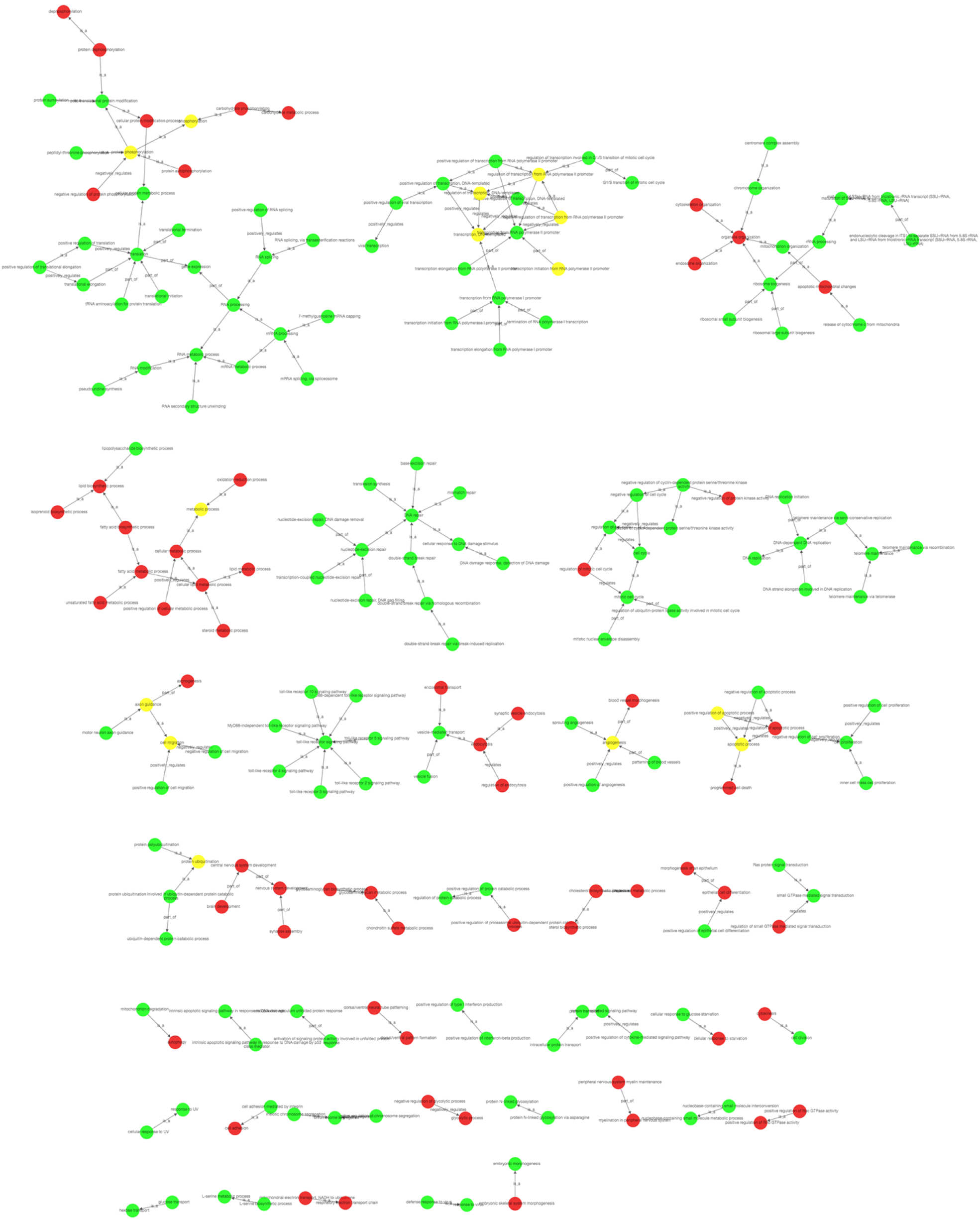
The GO-Tree constructed based on the GO and pathway analysis results. Red represents the significant GO terms or pathways of upregulated genes; Green represents the significant GO terms or pathways of downregulated genes; Yellow represents the significant GO terms or pathways of both upregulated and downregulated genes.

**Figure 6.**
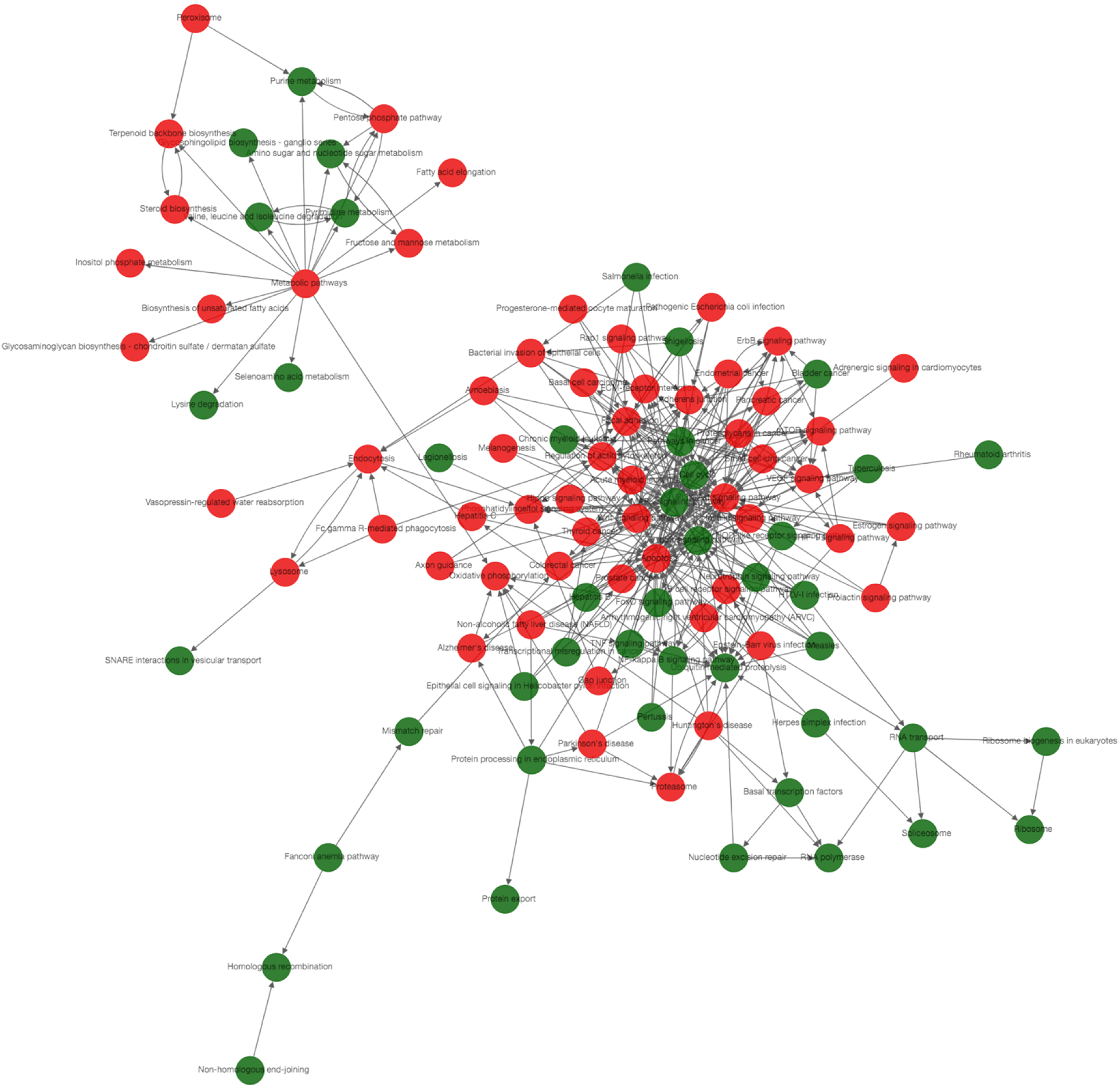
The Path-Act-Network constructed based on the GO and pathway analysis results. Red represents the significant GO terms or pathways of upregulated genes; Green represents the significant GO terms or pathways of downregulated genes.

### DEGs involved in PI3K-AKT signaling pathway

After analysis of RNA-seq data, the DEGs involved in PI3K-AKT signaling pathway were selected through combining with the KEGG database., PI3K and AKT were inevitable the hub nodes with highest degree in this signaling pathway. Besides, various signaling pathways, such as toll-like receptor signaling pathway, cell cycle, NF-KB signaling pathway and p53 signaling pathway were also regulated the expression of PI3K and AKT (Figure 7A). Furthermore, the DEGs in PI3K-AKT signaling pathway were clustered by heatmap. For instance, the up-regulated genes NOS3 and SGK3 and down-regulated genes FGF21 and VTN were clustered among of these DEGs (Figure 7B). In addition, western blot assay was performed to verified the expression of DEGs selected in PI3K-AKT signaling pathway. The results showed that the expression of ANGPT2, CD19, COL4A3, FGF18, ITGB4, ITGB8, LAMA3, LAMC2, PPP2R2C and SYK were upregulated in SKOV3 cell lines compared to that in MCV152 cell lines (Figure 8A), while compared with MCV152 cell lines, the protein levels of AKT3, COL6A1, CSF3, FGF1, ITGA2, ITGA11, MYB, PCK2, PGF, PIK3AP1, SGK1, TLR4 and TP53 were decreased in SKOV3 cell lines (Figure 8B), which were consistent with the results of bioinformatics analysis.

**Figure 7.**
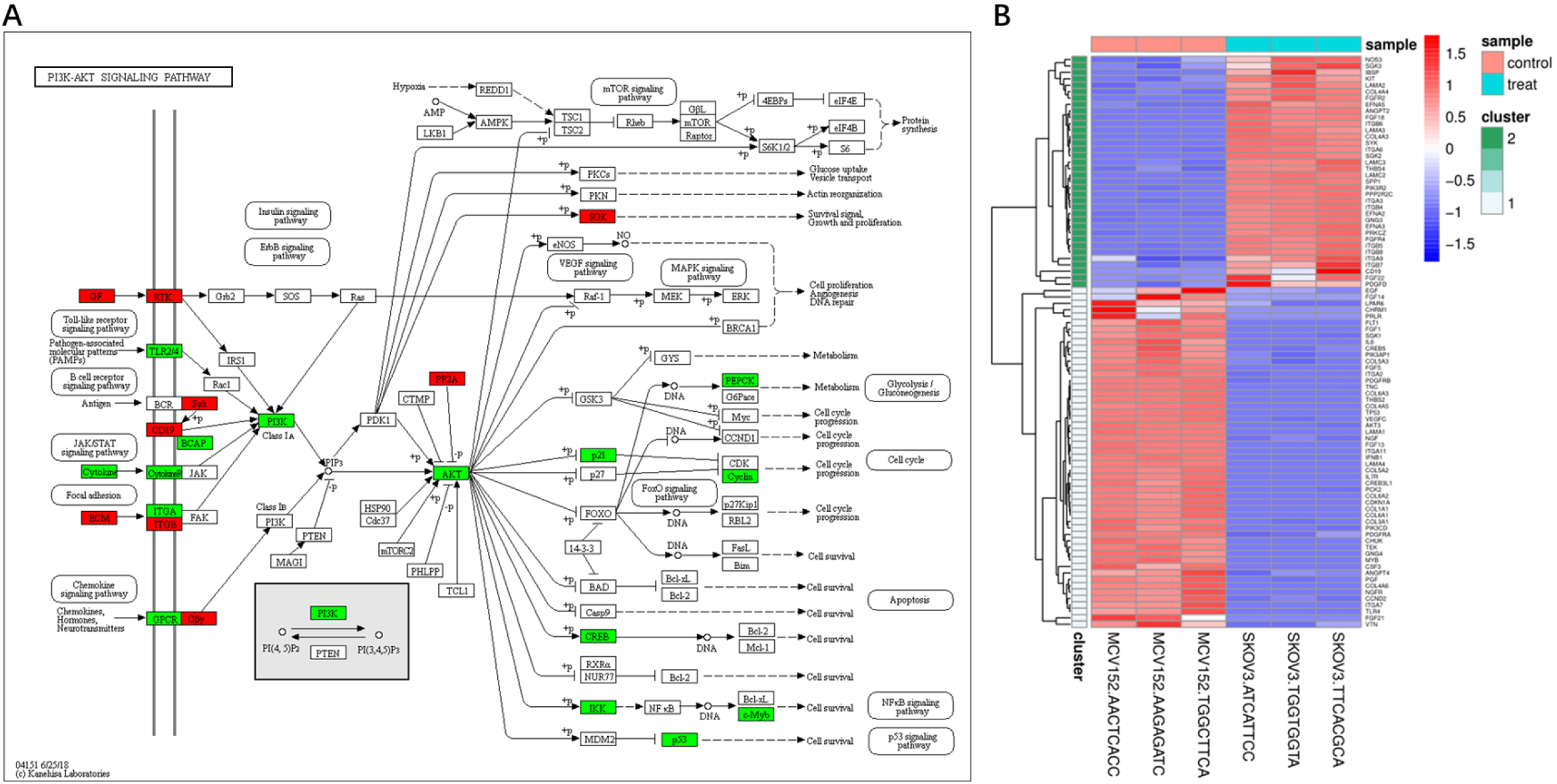
The DEGs involved in PI3K-AKT signaling pathway were selected through KEGG database and analyzed by heatmap. (A) The upregulated and down-regulated genes in PI3K-Akt signaling pathway. (B) Heatmap of genes involved in PI3K-Akt signaling pathway. X axis: sample name; Y axis: gene name. Red represents the significant upregulated genes; Green represents the significant downregulated genes.

**Figure 8.**
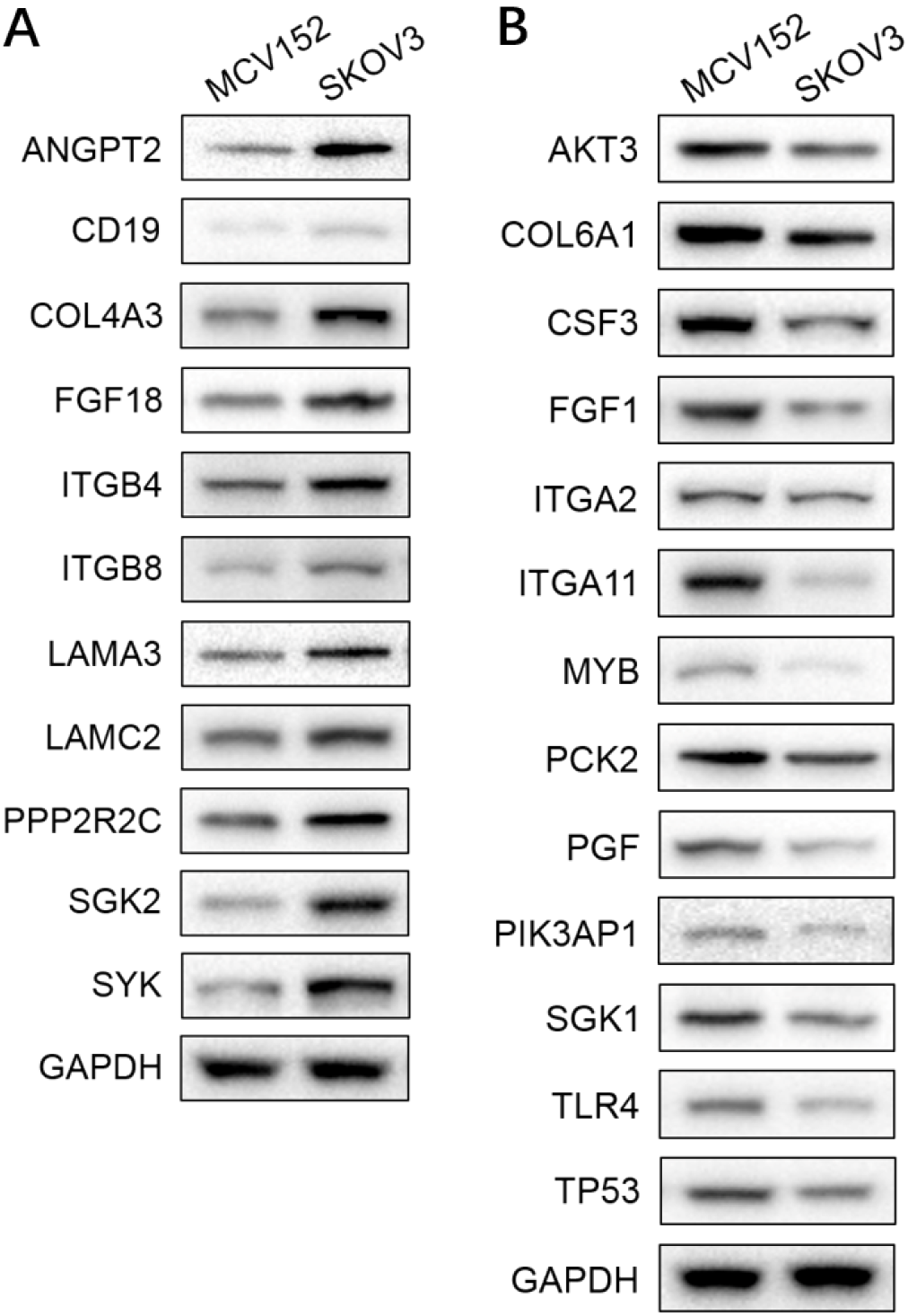
The protein levels of genes involved in PI3K-Akt signaling pathway detected by western blot. (A) The protein levels of ANGPT2, CD19, COL4A3, FGF18, ITGB4, ITGB8, LAMA3, LAMC2, PPP2R2C and SYK in MCV152 and SKOV3 cell lines. (B) The protein levels of AKT3, COL6A1, CSF3, FGF1, ITGA2, ITGA11, MYB, PCK2, PGF, PIK3AP1, SGK1, TLR4 and TP53 in MCV152 and SKOV3 cell lines.

## Discussion

In this study, 2,020 upregulated and 1,673 downregulated DEGs were obtained between SKOV3 and MCV152 cells. The upregulated and downregulated DEGs were significantly associated with cell adhesion. Furthermore, the upregulated DEGs were significantly involved in pathways of ECM-receptor interaction, and the downregulated DEGs were involved in PI3K-Akt signaling pathway. PCR and western blot analyses showed that the expression levels of genes (proteins) associated with PI3K-Akt signaling pathway were in consistent with bioinformatics analysis.

Functional analyses indicated that the upregulated DEGs were significantly correlated with cell adhesion and ECM-receptor interaction. Cell–cell adhesion is a fundamental biological process that defines morphogenesis of cells and tissues in multicellular organisms [13]. Overexpression of cell adhesion molecules, such as Claudin 3, DDR1 and Ep-Cam, were the early events in the development of ovarian cancer [14]. Besides, vascular cell adhesion molecule-1 was also a regulator of ovarian cancer peritoneal metastasis [15]. The signaling conveyed by cell adhesion and with the underlying ECM is closely coordinated with gene regulation during normal tissue homeostasis [16]. Disruption of cell adhesion, the subsequent changes in adhesion-mediated signal transduction, and the increase in cell motility are characteristics in the development of invasive metastatic cancer [17]. It has been reported that alterations in cell–cell and cell–ECM adhesion promote cancer cell invasion, intravasation and extravasation [18]. Therefore, cell adhesion and ECM-receptor interaction may play an important role in ovarian cancer invasion.

The PI3K-Akt signaling pathway regulates a variety of cellular processes including cell survival, proliferation, growth and motility, which are important for tumorigenesis. Inhibition of HER3-PI3K-AKT signaling pathway could enhance the apoptosis of ovarian cancer cells [19]. Mabuchi et al. also suggested that PI3K/AKT/mTOR pathway could be used as a therapeutic target for ovarian cancer [20]. Mutation or altered expression of components of this pathway is implicated in human cancer [21]. Importantly, this pathway was also enriched by DEGs in this study consistent with many other studies that had demonstrated a pivotal role for PI3K-Akt signaling pathway in ovarian cancer [20, 22, 23]. Based on the RNA-seq data and KEGG database, we validated the expression of DEGs and corresponding proteins associated with PI3K-Akt signaling pathway, such as the upregulated DEGs of *ANGPT2*, *FGF18*, *ITGB4* and *ITGB8*, and downregulated DEGs of *AKT3* and *PIK3AP1*.

ANGPT2 (angiopoietin 2) is a member of the human ANGPT-TIE system [24], which plays an important role in tumor initiation and increasing the number of tumor vessel sprouts [25, 26]. In some human cancers, overexpressed ANGPT2 is correlated with poor prognosis [27, 28]. FGF18 (fibroblast growth factor 18) is a member of the FGF family, acting as a chemotactic, mitogenic and angiogenic factor [29]. It has been suggested to play a critical role in promoting the progression of ovarian high-grade serous carcinoma [30]. ITGB4 (integrin beta-4) and ITGB8 are derived from integrin family which control cell attachment to the ECM and activate intracellular signaling pathways associated with cell survival, growth, differentiation, apoptosis, and migration [31]. Increased expression of ITGB4 and ITGB8 has been reported to be associated with tumorigenesis [32, 33]. Our results also suggested the increased expression of *ANGPT2*, *FGF18*, *ITGB4* and *ITGB8* in SKOV3 cells compared with that in MCV152 cells.

AKT3 (AKT serine/threonine kinase 3), a member of the AKT family, is the key regulator of many cellular processes associated with cancer, including cell proliferation, survival and metastasis [34]. Blockade of AKT3 results in a significant reduction in the expression of EBAG9, a tumor cell marker [35], whose expression is closely associated with VEGF expression in ovarian and uterine tumor samples [36]. PIK3AP1 (phosphoinositide-3-kinase adaptor protein 1) is identified as a tumor suppressor gene and has been reported to be inactivated in head and neck cancers [37]. But its role in ovarian cancer has not been fully investigated. In our study, the two genes were demonstrated to be significantly downregulated in SKOV3 cells compared with MCV152 cells, and both genes were involved in various signaling pathways, which suggested that downregulated AKT3 and PIK3AP1 may be associated with ovarian cancer development.

In conclusion, cell adhesion and ECM-receptor interaction may play an important role in ovarian cancer invasion. PI3K-Akt signaling pathway may involve in ovarian cancer progression by upregulating *ANGPT2*, *FGF18*, *ITGB4* and *ITGB8*, and downregulating *AKT3* and *PIK3AP1*.

## References

1. Reid, B.M., J.B. Permuth, and T.A. Sellers, Epidemiology of ovarian cancer: a review. Cancer biology & medicine, 2017. 14(1): p. 9.

2. Ferlay, J., et al., Cancer incidence and mortality worldwide: IARC CancerBase. GLOBOCAN 2012 v10, 2012. 11.

3. Howlader, N., et al., SEER cancer statistics review, 1975–2014. Bethesda, MD: National Cancer Institute, 2017. 2018.

4. Siegel, R.L., K.D. Miller, and A. Jemal, Cancer Statistics, 2017. Ca Cancer J Clin, 2017. 67(1): p. 7.

5. Karnezis, A.N., et al., The disparate origins of ovarian cancers: pathogenesis and prevention strategies. Nature Reviews Cancer, 2016. 17(1).

6. Wallace, S., et al., Efforts at maximal cytoreduction improve survival in ovarian cancer patients, even when complete gross resection is not feasible. Gynecologic oncology, 2017. 145(1): p. 21–26.

7. Quitadamo, A., et al., An integrated network of microRNA and gene expression in ovarian cancer. BMC Bioinformatics, 2015. 16(5): p. S5.

8. Welsh, J.B., et al., Analysis of gene expression profiles in normal and neoplastic ovarian tissue samples identifies candidate molecular markers of epithelial ovarian cancer. Proceedings of the National Academy of Sciences, 2001. 98(3): p. 1176–1181.

9. Nielsen, J.S., et al., Prognostic significance of p53, Her - 2, and EGFR overexpression in borderline and epithelial ovarian cancer. International Journal of Gynecological Cancer, 2004. 14(6): p. 1086–1096.

10. Santin, A.D., et al., Gene expression profiles in primary ovarian serous papillary tumors and normal ovarian epithelium: identification of candidate molecular markers for ovarian cancer diagnosis and therapy. International journal of cancer, 2004. 112(1): p. 14–25.

11. Arend, R.C., et al., The Wnt/β-catenin pathway in ovarian cancer: A review. Gynecologic oncology, 2013. 131(3): p. 772–779.

12. Cheaib, B., A. Auguste, and A. Leary, The PI3K/Akt/mTOR pathway in ovarian cancer: therapeutic opportunities and challenges. Chinese journal of cancer, 2015. 34(1): p. 4.

13. Wickstroem, S.A. and C.M. Niessen, Cell adhesion and mechanics as drivers of tissue organization and differentiation: local cues for large scale organization. Current opinion in cell biology, 2018. 54: p. 89–97.

14. Heinzelmann-Schwarz, V.A., et al., Overexpression of the cell adhesion molecules DDR1, Claudin 3, and Ep-CAM in metaplastic ovarian epithelium and ovarian cancer. Clinical Cancer Research An Official Journal of the American Association for Cancer Research, 2004. 10(13): p. 4427.

15. Slack-Davis, J., et al., Vascular Cell Adhesion Molecule-1 Is a Regulator of Ovarian Cancer Peritoneal Metastasis. Cancer Research, 2009. 69(4): p. 1469–1476.

16. Clément, R., et al., Viscoelastic dissipation stabilizes cell shape changes during tissue morphogenesis. Current Biology, 2017. 27(20): p. 3132–3142. e4.

17. Basu, S., S. Cheriyamundath, and A. Ben-Ze’ev, Cell–cell adhesion: linking Wnt/β-catenin signaling with partial EMT and stemness traits in tumorigenesis. F1000Research, 2018. 7.

18. Sousa, B., J. Pereira, and J. Paredes, The Crosstalk Between Cell Adhesion and Cancer Metabolism. International Journal of Molecular Sciences, 2019. 20(8): p. 1933.

19. Bezler, et al., Inhibition of doxorubicin-induced HER3-PI3K-AKT signalling enhances apoptosis of ovarian cancer cells. Molecular Oncology, 2012. 6(5): p. 516–529.

20. Mabuchi, S., et al., The PI3K/AKT/mTOR pathway as a therapeutic target in ovarian cancer. Gynecologic Oncology, 2015. 137(1): p. 173–179.

21. Vivanco, I. and C.L. Sawyers, The phosphatidylinositol 3-kinase–AKT pathway in human cancer. Nature Reviews Cancer, 2002. 2(7): p. 489.

22. Thant, A.A., et al., Fibronectin activates matrix metalloproteinase-9 secretion via the MEK1-MAPK and the PI3K-Akt pathways in ovarian cancer cells. Clinical & experimental metastasis, 2000. 18(5): p. 423–428.

23. Li, H., J. Zeng, and K. Shen, PI3K/AKT/mTOR signaling pathway as a therapeutic target for ovarian cancer. Archives of gynecology and obstetrics, 2014. 290(6): p. 1067–1078.

24. Huang, H., et al., Targeting the ANGPT–TIE2 pathway in malignancy. Nature Reviews Cancer, 2010. 10(8): p. 575.

25. Hashizume, H., et al., Complementary actions of inhibitors of angiopoietin-2 and VEGF on tumor angiogenesis and growth. Cancer research, 2010. 70(6): p. 2213–2223.

26. Nasarre, P., et al., Host-derived angiopoietin-2 affects early stages of tumor development and vessel maturation but is dispensable for later stages of tumor growth. Cancer research, 2009. 69(4): p. 1324–1333.

27. Sfiligoi, C., et al., Angiopoietin - 2 expression in breast cancer correlates with lymph node invasion and short survival. International journal of cancer, 2003. 103(4): p. 466–474.

28. J: son Lind, A., et al., Angiopoietin 2 expression is related to histological grade, vascular density, metastases, and outcome in prostate cancer. The Prostate, 2005. 62(4): p. 394–399.

29. Moore, E., et al., Fibroblast growth factor-18 stimulates chondrogenesis and cartilage repair in a rat model of injury-induced osteoarthritis. Osteoarthritis and cartilage, 2005. 13(7): p. 623–631.

30. S, E.-G., et al., FGF18 as a potential biomarker in serous and mucinous ovarian tumors. Tumour biology: the journal of the International Society for Oncodevelopmental Biology and Medicine, 2016. 37(3): p. 3173–83.

31. Hood, J.D. and D.A. Cheresh, Role of integrins in cell invasion and migration. Nature Reviews Cancer, 2002. 2(2): p. 91.

32. Xu, Z. and R. Wu, Alteration in metastasis potential and gene expression in human lung cancer cell lines by ITGB8 silencing. The Anatomical Record: Advances in Integrative Anatomy and Evolutionary Biology, 2012. 295(9): p. 1446–1454.

33. Knyazev, E., et al., Suppression of ITGB4 Gene Expression in PC-3 Cells with Short Interfering RNA Induces Changes in the Expression of [beta]-Integrins Associated with RGD-Receptors. Bulletin of experimental biology and medicine, 2015. 159(4): p. 541.

34. Bellacosa, A., S. Staal, and P. Tsichlis, A retroviral oncogene, akt, encoding a serine-threonine kinase containing an SH2-like region. Science, 1991. 254(5029): p. 274–277.

35. Chatterjee, M., et al., Diagnostic markers of ovarian cancer by high-throughput antigen cloning and detection on arrays. Cancer research, 2006. 66(2): p. 1181–1190.

36. Akahira, J., et al., Expression of EBAG9/RCAS1 is associated with advanced disease in human epithelial ovarian cancer. British journal of cancer, 2004. 90(11): p. 2197.

37. Guerrero-Preston, R., et al., Key tumor suppressor genes inactivated by “greater promoter” methylation and somatic mutations in head and neck cancer. Epigenetics, 2014. 9(7): p. 1031–1046.

